# Brain cell type nuclei enrichment without fixative for nanoCUT&Tag and other omics approaches

**DOI:** 10.64898/2026.03.31.715540

**Authors:** Kevin Chris Ziegler, Janna D. van Dalen, Lucy Bedwell, Janis Transfeld, Alexi Nott

## Abstract

We present a workflow for cell-type-enriched epigenomic profiling of the neurovascular unit and associated cell types, including brain endothelial cells, mural cells, microglia, astrocytes, neurons, and oligodendrocytes isolated from frozen unfixed human and mouse brain tissue. The workflow consists of an unfixed nuclei isolation with improved vascular nuclei release, fluorescence-activated nuclei sorting (FANS) based on cell-type-specific DNA-bound proteins, and nanoCUT&Tag for epigenomic profiling. The nanoCUT&Tag uses Tn5 transposase fused to a single-chain nanobody with secondary antibody-like properties. This allows low-input epigenomic profiling that is compatible with FANS-enriched nuclei immunolabeled for transcription factor markers, which is not possible with traditional CUT&Tag approaches. The protocol allows studying the human brain epigenome in a cell type-specific manner, which is increasingly associated with neurodegenerative diseases. The workflow can be used on various tissue sources, including resected and post-mortem archived brain tissue and can be coupled to multiple-omics approaches including single nuclei (sn)RNA-seq, snATAC-seq, and proteomics.

## Introduction

Multiple brain conditions, including Alzheimer’s disease (AD), schizophrenia, bipolar disorder, epilepsy, and autism spectrum, often develop sporadically and are partly caused by a combination of risk variants with small effect sizes^1,2^. Genome-wide association studies (GWAS) have identified thousands of risk variants for these conditions, which are mostly found in non-coding regions^3^. These non-coding variants influence cis-regulatory elements (CREs), such as enhancers, promoters, and silencers^4^. Enhancers are highly cell-type-specific regulatory regions that drive gene expression of distal genes and were identified as major carriers of genetic risk^5,6^.

Microglia are brain resident immune cells and have been associated with the genetic risk of AD^6–8^ and other neurodegenerative conditions^9,10^. Microglia constitute ∼5% of the total cells within the brain parenchyma, highlighting the importance of cell type enrichment strategies over bulk approaches. Recent single-cell gene expression studies have shown that many of the risk genes identified in microglia are similarly expressed in vascular brain endothelial cells^11,12^. However, vascular cell types can be difficult to isolate from the vessel basement membrane. We developed a non-fixed nuclei isolation approach followed by nanobody-based cleavage under targets and tagmentation (nanoCUT&Tag) that allowed epigenomic profiling of brain cell types, including endothelial cells and mural cells^13,14^. Using this approach, we showed that AD heritability is strongly associated with microglia/myeloid cells, with some enrichment for vascular cell types^13^. Whereas small vessel disease, which is often comorbid with AD, was strongly associated with the major brain blood vessel cell types, endothelial cells, mural cells and astrocytes^13^. The cell type risk genes we identified were mostly disease-specific, indicating unique vascular and immune mechanisms across these pathologies^13^. Lastly, we integrated gene expression analysis to prioritise repurposable drugs with genetic evidence for the treatment of AD^13^.

### Overview of the technique

Rare cell types, such as those of the neurovascular unit, are difficult to isolate and profile using standard methods, such as chromatin immunoprecipitation followed by sequencing (ChIP-seq)^15^, due to the need for a high number of cells. To address this, we developed a workflow that increases the yield of vascular nuclei and lowers nuclei input requirements for epigenomic profiling (**Fig. 1**)^13^. This workflow consists of nuclei isolation from fresh-frozen human^13^ or mouse brain tissue^14^. Brain tissue (100-500 mg) is minced into ∼1 mm pieces and subsequently Dounce homogenised. The homogenate is gently mashed through strainers to dissociate nuclei retained in remaining blood vessels, followed by a dextran clean-up step, which increases nuclei purity, improving sorting time and efficiency during FANS. The sample is stained overnight with antibodies against cell-type-specific nuclear markers: ERG for endothelial cells, NOTCH3 for human mural cells (pericytes and smooth muscle cells), SUR2B for mouse mural cells, RFX4 for human astrocytes, LHX2 for mouse/human astrocytes, PU1 for microglia, NEUN for neurons, and OLIG2 for oligodendrocytes.

**Figure 1:**
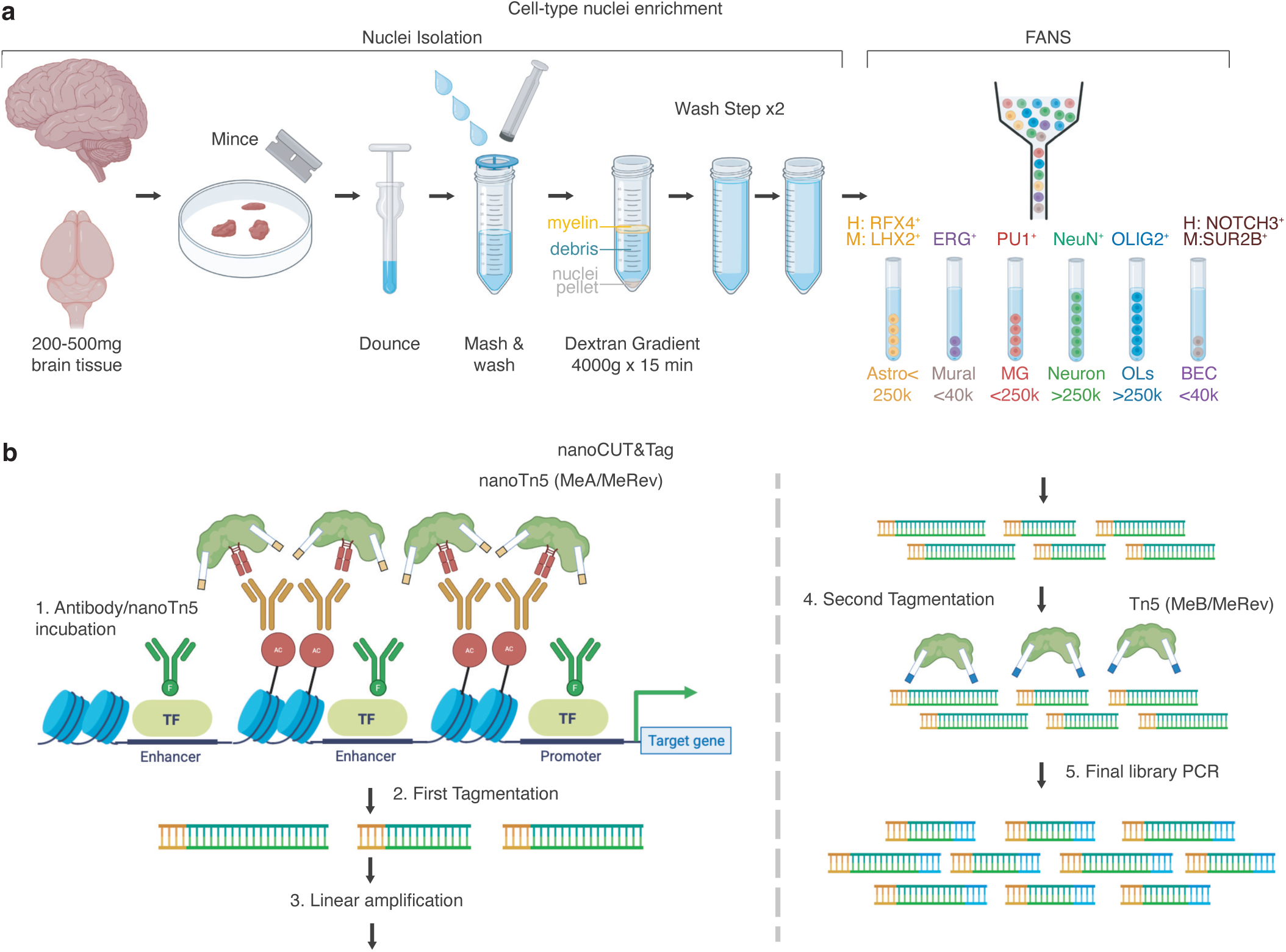
Workflow for brain cell-type-enriched epigenomic profiling. **(A)** Workflow for unfixed nuclei isolation and cell type enrichment using FANS. Tissue is minced using a razor blade, Dounce homogenised, and vascular nuclei released by mashing. Debris and myelin are removed using an 18% dextran centrifugation. Cell type nuclei are enriched by FANS using ERG/FLI1 (brain endothelial cells), NOTCH3 (human mural cells), SUR2B (mouse mural cells), PU1 (microglia/myeloid cells), NeuN (neurons), OLIG2 (oligodendrocytes), and LHX2 (mouse/human astrocytes), RFX4 (human astrocytes). **(B)** Workflow for bulk nanoCUT&Tag with linear amplification. The first nanoTn5 tagmentation recognises species-specific immunoglobulins and is designed to avoid FANS-antibody binding (steps 1 & 2). A linear PCR introduces the i5 adapters (step 3), followed by a second tagmentation (step 4) and exponential PCR (step 5) to introduce i7 adapters for final library preparation.

The cell-type-specific nuclei are sorted by FANS, and a maximum of 50,000 nuclei are used for epigenomic profiling using a modified version of nanoCUT&Tag (**Fig. 1**). Histone modifications or transcription factors (TFs) are targeted using a primary antibody, followed by recruitment of Tn5 linked to a single-domain nanobody that functions as a secondary antibody to recognize host rabbit or host mouse primary antibodies. The nano-Tn5 tagments the chromatin to produce target-specific DNA fragments with inserted adapters. DNA fragments are amplified using a linear single-primer polymerase chain reaction (PCR), followed by a second tagmentation using a Tn5 tagmentase. Final libraries are amplified by exponential PCR, quantified using Qubit, and fragment size distribution determined using a TapeStation. The libraries undergo next-generation sequencing using Illumina technology and are analysed using Nextflow pipelines.

### Applications of the method

The non-fixed nuclei isolation protocol can be used to profile rare or difficult to isolate cell types using nanoCUT&Tag^13^ (**Fig. 1**). This approach can enrich for nuclei from parenchymal and vascular brain compartments. This allows for a more complete analysis of the neurovascular unit and can be adapted for additional cell types and states upon the development of new nuclear markers.

Our non-fixed nuclei enrichment protocol is coupled with a nanoCUT&Tag workflow for downstream profiling of histone modifications and chromatin factors in cell-type-enriched nuclei^13^ (**Fig. 1**). This approach has been validated in human and mouse brain for histone H3 lysine 27 acetylation (H3K27ac) and H3 lysine 4 trimethylation (H3K4me3) (**Fig. 2, 3**)^13^, which are markers of enhancers and promoters respectively, as well as repressive H3K27me3. Transcription factors that can be profiled using nanoCUT&Tag include PU1, ERG/FLI1 and OLIG2 in microglia, brain endothelial cells and oligodendrocytes, respectively, although this is dependent on transcription factor affinity and antibody available.

**Figure 2:**
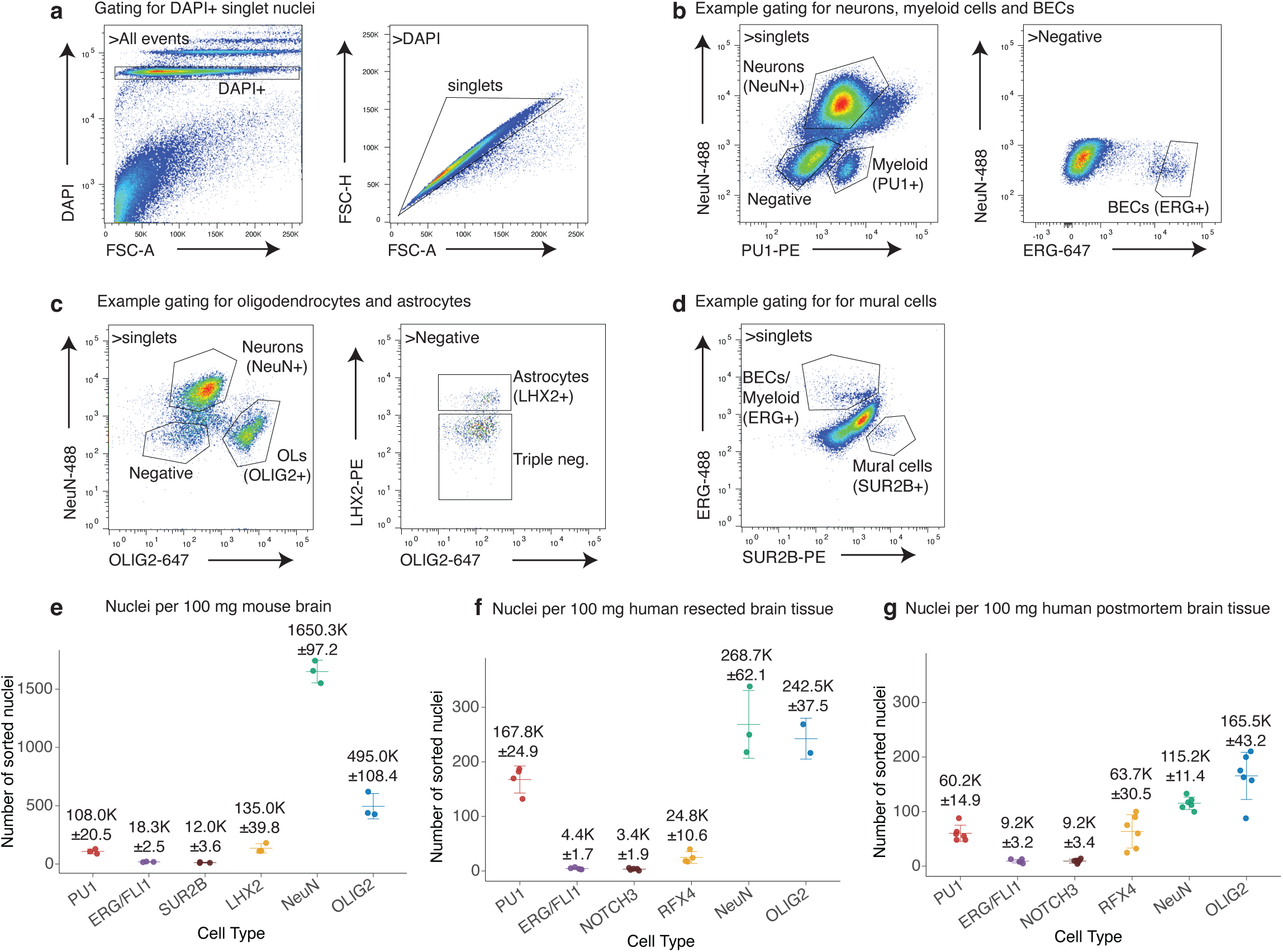
FANS gating strategy for brain vascular and non-vascular cell types. **(A)** Gating to select mouse DAPI+ nuclei (DAPI vs. FSC-A) and exclude doublets (FSC-H vs. FSC-A). **(B)** Mouse nuclei gating to enrich for neurons (NeuN^+ve^, PU1^-ve^), microglia/myeloid cells (NeuN^-ve^/PU1^+ve^), and brain endothelial cells (NeuN^-ve^/PU1^-ve^/ERG^high^) using NeuN-488, PU1-PE, and ERG-647. **(C)** Mouse nuclei gating to enrich for neurons (NeuN^+ve^/OLIG2^-ve^), oligodendrocytes (NeuN^-ve^/OLIG2^+ve^), and astrocytes (NeuN^-ve^/OLIG2^-ve^/LHX2^+ve^) using NeuN-488, OLIG2-647, and LHX2-PE. In human tissue, LHX2-PE can be replaced with RFX4-PE. **(D)** Mouse nuclei gating to enrich for mural cells (ERG^-ve^/SUR2B^+ve^) using SUR2B-PE and ERG-488. In human tissue, NOTCH3-PE is used to enrich for mural cells. **(E)** Expected nuclei numbers for ∼100 mg mouse brain tissue for each cell type. **(F)** Expected nuclei numbers for ∼100 mg human resected brain tissue for each cell type. **(G)** Expected nuclei numbers for ∼100 mg human postmortem brain tissue for each cell type. Bar plots represent mean values and standard deviations.

**Figure 3:**
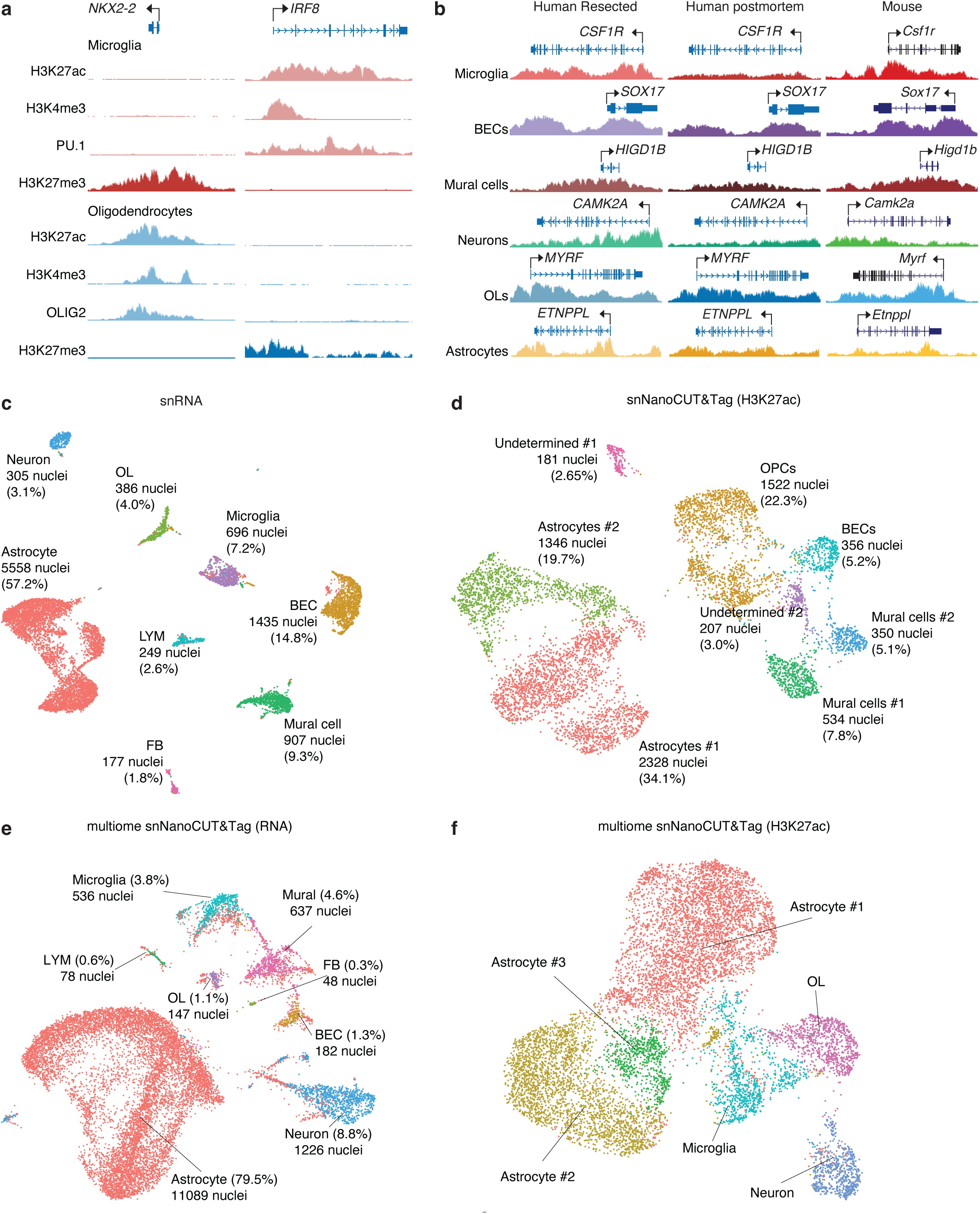
Workflow for brain cell-type-enriched epigenomic profiling. **(A)** Workflow for unfixed nuclei isolation and cell type enrichment using FANS. Tissue is minced using a razor blade, Dounce homogenised, and vascular nuclei released by mashing. Debris and myelin are removed using an 18% dextran centrifugation. Cell type nuclei are enriched by FANS using ERG/FLI1 (brain endothelial cells), NOTCH3 (human mural cells), SUR2B (mouse mural cells), PU1 (microglia/myeloid cells), NeuN (neurons), OLIG2 (oligodendrocytes), and LHX2 (mouse/human astrocytes), RFX4 (human astrocytes). **(B)** Workflow for bulk nanoCUT&Tag with linear amplification. The first nanoTn5 tagmentation recognises species-specific immunoglobulins and is designed to avoid FANS-antibody binding (steps 1 & 2). A linear PCR introduces the i5 adapters (step 3), followed by a second tagmentation (step 4) and exponential PCR (step 5) to introduce i7 adapters for final library preparation.

This workflow can be adapted for other modalities that require unfixed material, such as RNA-seq and ATAC-seq (including single cell), and, as recently shown, for proteomics of nuclear proteins^14^. The nuclei isolation protocol is compatible with postmortem tissue, enabling translational analysis of archived human specimens. In summary, FANS coupled with nanoCUT&Tag represents a powerful and versatile tool for investigating the epigenomic landscapes of specific cell types in the brain.

### Comparison with other methods

The fixed nuclei isolation protocol^16^ is optimized for ChIP-seq and has generated high-quality reference maps for the major brain cell types^6^. However, ChIP-seq requires >0.5 M nuclei and fixed nuclei are not compatible with other approaches sensitive to tissue fixation^6,16^. The non-fixed nuclei isolation allows for CUT&Tag profiling of histone modifications and some transcription factors for low input and single cell analysis. Traditional CUT&Tag protocols use Tn5 transposase coupled to Protein A (pA), which binds the DNA-binding transcription factors used to FANS-enrich nuclei, generating transcription factor CUT&Tag profiles. For analysis of histone modifications and other chromatin factors, we adapted a nanoCUT&Tag protocol^17,18^, which uses a nanobody-coupled Tn5 that detects species-specific immunoglobulins. Implementation of nanoCUT&Tag allowed for specific profiling of histone modifications without contaminating CUT&Tag signal from transcription factors used to FANS-enrich nuclei. This approach can be adapted to generate multimodal nanoCUT&Tag data of histone modifications together with transcription factor profiling from the same sample^17^.

Protocols for isolating non-fixed nuclei from archived brain tissue have allowed omics analysis of the major brain cell types^19,20^. However, these do not enrich for brain endothelial cells and mural cells that are difficult to release from blood vessels, and for mural cells there was a lack of validated nuclear markers. Optimisation of the non-fixed nuclei isolation, incorporating vascular nuclei-release steps adapted from VINE-seq^12,21^, increased the yield of brain endothelial nuclei. To capture mural cells, nuclei were stained for notch receptor 3 (NOTCH3), which regulates transcription within the nucleus upon ligand activation. Astrocytes, a major cellular component of the neurovasculature, can be enriched using LHX2^6,16^, though this marker may not capture the full diversity of astrocytic subtypes. We examined a new nuclear marker for astrocytes, RFX4, that broadly captures the heterogeneity of astrocytes in the cortex^13^. In summary, the non-fixed isolation of cell type nuclei coupled to nanoCUT&Tag allows for low input profiling of nuclei from difficult to isolate cell types.

## Experimental design

### Biological replicates

FANS coupled with nanoCUT&Tag was optimised for the major brain cell types, generates high-quality data across species (human and mouse), and is robust to different brain tissue acquisition methods, including resected and post-mortem samples. For nanoCUT&Tag profiling of histone modifications, we recommend using 5,000-50,000 nuclei per sample. For comparisons across different cell types, 3-4 biological replicates per group capture most cell-type-specific features^6,13^. In contrast, studies comparing two conditions within a single cell type, such as health versus disease, require sample sizes of 20 or more per group, although prior studies reported similar comparisons using ∼10 samples per group^22^.

### Tissue preparation and dissociation

Unfixed human brain tissue should be handled in a biosafety cabinet, and all procedures performed on ice. Repeated freeze-thaw cycles adversely affect nuclei integrity and the stability of DNA-bound proteins, including histones. To minimise tissue degradation, brain samples should be aliquoted in a single session on dry ice into ∼250 mg aliquots. This tissue quantity yields sufficient nuclei (>5,000) for difficult-to-isolate cell types, such as brain endothelial and mural cells. On the day of the experiment, the tissue is thawed in ice-cold lysis buffer and finely minced with a razor blade into ∼1 mm fragments. The minced tissue is Dounce homogenized using a loose pestle, followed by a tight pestle, and then passed through a 70 µm cell strainer.

Vessel fragments retained on the strainer are released by gentle mashing with the rubber plunger of a syringe and washing with Nuclei Wash Buffer. This procedure is repeated using a 40 µm strainer, through which the entire tissue suspension is filtered. Following an initial centrifugation step, the supernatant is discarded and the pellet resuspended. To efficiently remove debris and myelin, a wash and centrifugation step in 18% dextran is performed, followed by a wash and centrifugation in Nuclei Wash Buffer. The final nuclei pellet is resuspended in FANS Buffer.

### Cell type nuclei isolation by FANS

Nuclei were stained using a panel of antibodies targeting nuclear proteins enriched in specific brain cell types. These included NeuN for neurons, PU1 for microglia/myeloid cells, OLIG2 for oligodendrocytes, LHX2 or RFX4 for astrocytes, ERG/FLI1 for endothelial cells, and SUR2B (mouse) or NOTCH3 (human) for mural cells. Nuclei were enriched using a flow cytometer sorter (BD FACSMelody and BD FACSAria II), supporting three to four fluorophores. When modifying the antibody panel, users should evaluate spectral overlaps, include appropriate isotype controls, consider instrument-specific parameters and perform spectral compensation.

For several cell types, combinatorial gating strategies are required. LHX2 is expressed in neurons and astrocytes, and requires co-staining with NeuN to distinguish astrocytes (NeuN^-ve^, LHX2^+ve^). Alternatively, RFX4 labels most astrocytes with low-level expression in other cell types, and should ideally be co-stained with NeuN, OLIG2, and PU1. Similarly, a subset of microglia/myeloid cells are positive for ERG/FLI1, and additional myeloid populations may express low levels of both ERG/FLI1 and PU1. Therefore, nuclei for brain endothelial cells were enriched as an ERG^high^/PU1^-ve^ population.

Overnight immunostaining at 4 °C is recommended; however, shorter incubation times may be sufficient for some antibodies, such as NeuN and OLIG2. Antibodies that are unconjugated (LHX2, RFX4, and SUR2B) require a secondary antibody incubation step, which is performed the following day for 1 hour at 4 °C. Prior to sorting, stained nuclei are filtered through a 35 µm strainer and diluted to a final concentration of ∼1 million nuclei/mL to prevent clogging the sorter. This concentration supports high-purity sorting at an event rate of 1,000–5,000 events/s. Sort nozzles of 85 µm or 100 µm are recommended. Between 5,000 and 50,000 nuclei are typically collected into 1.5 or 2 mL tubes. Excess nuclei may be collected and cryopreserved in Bambanker at −80 °C.

### Bulk nanoCUT&Tag

For epigenetic profiling using nanoCUT&Tag, it is recommended to proceed directly after sorting nuclei. NanoTn5 enables direct recruitment of the transposase to the target histone modification or transcription factor, without binding to the antibody used for FANS. The nanoTn5 can be produced in-house or by protein production facilities, and commercial versions are becoming available. The antibody for the target histone modification or transcription factor must be generated in a host species different from that used for FANS.

For nanoCUT&Tag, cell-type FANS-enriched nuclei are immobilised on activated Concanavalin A magnetic beads, allowing efficient washing and exchange of reagents. Nuclei are incubated with primary antibodies for the target histone modification or transcription factor. Unbound antibody is removed, and nanoTn5 loaded with the MeA/MeRev adapter is added using nanoTn5 that matches the host species of the primary antibody. Unbound nanoTn5 is removed and tagmentation is initiated, resulting in site-specific MeA adapter insertion. The tagmentation is terminated and DNA is released for linear PCR-based addition of the i5 barcode into each fragment.

Linear amplification ensures that fragments generated by single-adapter insertions are pre-amplified and retained for library construction, improving sensitivity and data quality. Next, Tn5 tagmentase loaded with the MeB/MeRev adapter randomly fragments the linear-amplified DNA, generating uniform fragment size distributions. This mitigates polymerase bias towards shorter fragments during the final exponential PCR amplification that introduces the i7 barcode. Final libraries are sequenced at ∼15 million paired-end reads per sample, which provides robust coverage for downstream analyses.

### Other downstream applications

The unfixed FANS protocol can be adapted to a negative sorting strategy, to enable unbiased enrichment of brain vascular cell populations. Nuclei can be stained with antibodies against NeuN, OLIG2, and PU1 to label neurons, oligodendrocytes, and myeloid cells, respectively. This allows sorting of a triple-negative population, which enriches for astrocytes and vascular cell types. Additional inclusion of LHX2 or RFX4 in the staining panel allows sorting of a quadruple-negative population, which further enriches for rare brain endothelial cells and mural cells.

Single-nuclei RNA-sequencing (sn)RNA-seq, snATAC-seq and snNanoCUT&Tag have been performed using unfixed FANS-enriched nuclei without additional protocol modifications. The snNanoCUT&Tag was successful for neurons, oligodendrocytes, astrocytes and microglia, whereas vascular cell types were prone to nuclei clumping following snNanoCUT&Tag. Nuclei clumping can be mitigated by reducing the number of washes and/or the omission of the EDTA-based tagmentation stop step.

The unfixed nuclei isolation protocol can be coupled to the 10x Genomics Multiome (RNA+ATAC) workflow, which simultaneously profiles gene expression and chromatin accessibility. To preserve RNA integrity, unfixed nuclei were stained for 3 hours, and FANS was performed on the same day. Immediately following sorting, nuclei were simultaneously incubated with primary histone antibodies and nanoTn5 for 3 hours. For compatibility with the 10X Genomics Multiome workflow, nanoTn5 was loaded with both MeA/MeRev and MeB/MeRev adapters, followed by a single tagmentation step without linear amplification. Nuclei tagmentation was completed on the same day and then loaded on to Chromium Next GEM Chip for partitioning into droplets containing uniquely barcoded GEM beads. Downstream processing followed the manufacturer’s user guide.

### Expertise needed to implement the protocol

Biosafety and tissue handling should be considered when working with unfixed human tissue, which can harbour pathogens transmissible through cuts or aerosols during centrifugation and FANS. Proper biosafety training should be mandatory, and all work should be performed in certified biosafety cabinets. Knowledge of institutional ethics, patient consent, and appropriate waste disposal is recommended. Brain tissue is delicate and requires careful handling, including control of temperature and physical force during Dounce homogenisation and mashing steps. Assessment of nuclei integrity and debris contamination by microscopy is recommended for quality control.

FACS/FANS expertise is required for isolating cell-type-enriched nuclei populations and is often available through dedicated facilities. This includes designing staining panels, applying appropriate isotype and compensation controls, and correctly implementing positive, negative, or dumping channel strategies. Operators should be able to troubleshoot event rates, nozzle clogging, and gating strategies to reliably enrich target populations.

Prior familiarisation of the CUT&Tag and nanoCUT&Tag protocols is recommended, including handling of nuclei and sensitive reagents such as antibodies and enzymes. While not essential, knowledge of low-input sample handling, adapter design, linear and exponential PCR amplification, and tagmentation is beneficial. For single-cell applications, timing-sensitive steps should be well coordinated, such as nuclei staining, FANS, tagmentation, operation of the Chromium X controller, as well as inclusion of RNA preservation for Multiome and snRNA-seq experiments. Knowledge of DNA library quality controls and sequencing requirements is recommended and can be provided by dedicated sequencing facilities. This includes measuring DNA concentration, evaluating fragment size distributions, and planning appropriate sequencing depth.

### Limitations

The protocol has been tested exclusively in brain tissue. The protocol is expected to work for other tissues; however, optimisation may be required for tissue dissociation and FANS for non-brain cell populations. The unfixed nuclei isolation was optimized to facilitate the release of vascular nuclei from the vascular basement membrane. However, the release of vascular nuclei is not fully efficient and may require enzymatic treatment. It remains unclear whether certain vascular subtypes are preferentially released during mechanical dissociation. Similarly, rare or fragile cell populations may provide very few nuclei, limiting library complexity and downstream data quality. Post-mortem interval and tissue handling can further affect nuclear membrane integrity.

The protocol has been described for enriching the major broad cell-types of the brain. However, this could be adapted to capture cell subtypes and substates, for example, excitatory and inhibitory neuronal subtypes or mural subtypes (pericytes and smooth muscle cells). This would require the optimisation of additional nuclear markers. In complement, single-cell approaches could capture finer cellular heterogeneity. Potential biases of the current antibodies toward specific subpopulations remain unknown, and for some cell types, such as fibroblasts or ependymal cells, suitable FANS antibodies were not tested. The limited fluorophores available for nuclear markers restricts simultaneous detection of multiple cell types, which can impact the isolation of rare subpopulations by FANS.

nanoCUT&Tag uses *in situ* enzymatic fragmentation whereas ChIP-seq uses sonicated crosslinked soluble chromatin, which may bias the regions profiled by each technique. Tagmentation efficiency depends on the DNA sequence and chromatin context, which may lead to uneven representation across the genome. Consequently, quantitative interpretation of read coverage as a measure of nucleosome occupancy is challenging, as signal strength can be influenced by antibody efficiency, chromatin accessibility, and tagmentation bias. The field lacks a definitive ground truth for chromatin states, and orthogonal validation experiments using independent assays are recommended.

Single-cell assays are constrained by nuclei clumping following FANS, especially for vascular cell types, which can be impacted by protocol length and mechanical stress. Next-generation sorters, such as the MACSQuant® Tyto®, may help reduce stress through low-pressure microfluidics. RNA integrity may be impacted for the 10X Genomics Multiome assays following unfixed FANS. Shortening the protocol to preserve RNA integrity can be optimised to avoid compromising the quality of epigenetic data. Rare populations may be underrepresented due to loss during FANS or nuclei clumping, and batch effects can introduce additional variability.

Overall, the protocol provides a robust framework for epigenetic profiling of brain cell types. However, further optimisation may be needed for tissue generalizability, subpopulation profiling, nuclei yield, antibody specificity, technical biases, and single-cell recovery.

## Materials

### Biological materials

Protocol optimisation and validation were performed using mouse and human brain tissue. Whole mouse brains from C57BL/6 mice were provided by Charles River UK Ltd. Resected temporal, parietal, and frontal cortical samples were obtained from paediatric epilepsy patients and snap-frozen immediately after surgery, as published^13^. Human post-mortem prefrontal cortical tissue (BA9) was obtained from aged individuals without cognitive impairment, as published^13^. Resected brain tissue was collected with informed consent under a protocol approved by the Institutional Review Board (IRB #171361) of UC San Diego and Rady Children’s Hospital and metadata reported^13^. Informed consent was obtained from the parents of all study participants, and when applicable, assent was also obtained from the patients. Fresh frozen post-mortem prefrontal cortex was obtained from Banner Sun Health Research Institute Brain and Body Donation Program of Sun City, Arizona and metadata reported^13^. All human tissue was stored at −80 °C until nuclei isolation, as published^13^.

### Reagents

- Protease Inhibitor Cocktail (Sigma-Aldrich, Cat # P8340)
- Proteinase K (New England Biolabs, Cat # P8107S)
- SPRIselect DNA Size Selection Reagent (Beckman Coulter, Cat # B23318)
- BioMagPlus® Concanavalin A beads (Bangs Laboratories, Cat # BP531-BAN)
- Tagmentase (Tn5 transposase) – unloaded (Diagenode, Cat # C01070010)
- NEBNext® Ultra™ II Q5® Master Mix (New England Biolabs, Cat # M0544L)
- Digitonin (Sigma-Aldrich, Cat # D5628-1G)
- Sucrose (VWR Life Science, Cat # 0335-500G)
- Calcium chloride (Sigma-Aldrich, Cat # C1016)
- Magnesium acetate solution (Sigma-Aldrich, Cat # 63052)
- UltraPure™ 0.5 M EDTA, pH 8.0 (Thermo Fisher Scientific, Cat # 15575020)
- UltraPure™ 1 M Tris-HCI, pH 8.0 (Thermo Fisher Scientific, Cat # 15568025)
- Triton™ X-100 (Sigma-Aldrich, Cat # T8787)
- Dithiothreitol (Thermo Fisher Scientific, Cat # BP172-5)
- Spermidine (Sigma-Aldrich, Cat # S2626-1G)
- HEPES, 1 M, pH 7.2 (Rockland, Cat # MB-062-0100)
- HEPES, 1 M, pH 7.5 (Thermo Fisher Scientific, Cat # J60712.AK)
- HEPES, 1 M, pH 8.0 (Thermo Fisher Scientific, Cat # J63578.AK)
- Ethanol, Molecular Biology Grade (Thermo Fisher Scientific, Cat # 16685992)
- Manganese(II) chloride tetrahydrate (Sigma-Aldrich, Cat # 203734-5G)
- Bovine Serum Albumin (Sigma-Aldrich, Cat # A3059-100G)
- Sodium chloride, 5 M (Thermo Fisher Scientific, Cat # AM9759)
- IGEPAL® CA-630 (NP-40) (Sigma-Aldrich, Cat # I8896-50ML)
- Potassium chloride, 2 M (Thermo Fisher Scientific, Thermo Fisher Scientific)
- Dulbecco’s Phosphate-Buffered Saline (DPBS) no calcium, no magnesium (Thermo Fisher Scientific, Cat # 14190144)
- Magnesium chloride, 1 M (Thermo Fisher Scientific, Cat # AM9530G)
- UltraPure™ SDS Solution, 10% (Thermo Fisher Scientific, Cat # 24730020)
- UltraPure™ Distilled Water (Thermo Fisher Scientific, Cat # 10977035)
- UltraPure™ Glycerol (Thermo Fisher Scientific, Cat # 15514011)
- ChIP DNA Clean & Concentrator Kit (Zymo, Cat # D5205)
- Qubit™ 1X dsDNA Assay-Kit (Invitrogen, Cat # 15860210)
- High Sensitivity DNA Kit (Agilent, Cat # 5067-4626)

### Oligonucleotides

Tn5 adapter sequences (taken from Kaya-Okur et al. 2020^23^)

- MeA: TCGTCGGCAGCGTCAGATGTGTATAAGAGACAG
- MeB: GTCTCGTGGGCTCGGAGATGTGTATAAGAGACAG
- MeRev: [PHO]CTGTCTCTTATACACATCT

Indexing PCR primers were taken from Buenrostro et al., 2015^24^

### Equipment

- 7 mL Wheaton^TM^ Dounce Tissue Grinder, Thermo Fisher Scientific, Cat # 06435A
- Falcon® 70 µm Cell Strainer, Corning, Cat # 352350
- Falcon® 40 µm Cell Strainer, Corning, Cat # 352340
- Qubit™ Assay Tubes, Thermo Fisher Scientific, Cat # Q32856
- 0.2 ml 8-Strip PCR Tubes, Individually Attached Domed Caps, Starlab, Cat # I1402-2900
- Eppendorf® DNA LoBind® 1.5 mL, VWR Life Science, Cat # 525-0130
- Eppendorf® DNA LoBind® 2.0 mL, VWR Life Science, Cat # 525-0131
- Fisherbrand™ Sterile Syringes for Single Use (5 mL), Thermo Fisher Scientific, Cat # 15809152
- Azpack™ Carbon Steel Razor Blades, Thermo Fisher Scientific, Cat # 11904325
- NEBNext® Magnetic Separation Rack, New England Biolabs, Cat # S1515S
- Falcon 5mL Round Bottom Polystyrene Test Tube with Cell Strainer Cap, Scientific Laboratory Supplies, Cat # 352235
- Falcon 5mL Round Bottom Polypropylene Test Tube with Snap Cap, Scientific Laboratory Supplies, Cat # 352063
- Petri Dish, VWR Life Science, Cat # 391-0467
- Screw cap tube, 15 ml, Sarstedt, Cat # 62.554.502
- Screw cap tube, 50 ml, Sarstedt, Cat # 62.547.254

### Reagent Setup

#### 0.1 M Dithiothreitol (DTT)

Dilute 0.1 mL 1 M DTT in 0.9 mL nuclease-free water. Store at −20°C for up to 1 month.

! CAUTION DTT solutions are harmful if swallowed and cause skin and serious eye irritation. May cause mild respiratory irritation if aerosolised. Handle with gloves and eye protection. Avoid splashing or creating aerosols. Work in a well-ventilated area or fume hood and wash hands after handling.

#### 10% (vol/vol) Triton X-100 stock solution

Dilute 1 mL Triton X-100 in 9 mL nuclease-free water in a 15 mL tube and mix for 30 min on an orbital shaker. Store protected from light at 4°C for up to 6 months.

! CAUTION Triton X-100 solutions cause skin and eye irritation and may cause respiratory irritation if aerosolised. Harmful to aquatic life. Handle with gloves and eye protection and avoid splashing or generating aerosols. Work in a well-ventilated area or fume hood and wash hands thoroughly after handling.

#### 20% Bovine Serum Albumin (BSA)

Dissolve 2 g BSA powder in 5 mL nuclease-free water. Adjust final volume to 10 mL with nuclease-free water. Sterile filter and store at 4°C for no longer than 1 week.

#### Nuclei Wash Stock

Dissolve 54.8 g sucrose (MW: 342.3 g mol^-1^) in 250 mL nuclease-free water. Add 2.5 mL 1 M CaCl_2_, 1.5 mL Mg-Acetate, 1 mL 0.5 M EDTA and 5 mL 1 M Tris-HCL pH 8.0. Adjust final volume to 500 mL with nuclease-free water. Sterile filter and keep at 4°C for up to 6 months.

#### Nuclei Wash Buffer

Add 200 µL protease inhibitor cocktail (PIC) (100x) and 1 mL 20% BSA to 18.8 mL Nuclei Wash Stock.

#### Lysis Buffer

Add 50 µL 10% (vol/vol) Triton X-100 stock solution, 50 µL of 0.1 M DTT and 50 µL PIC (100X) to 4.85 mL of Nuclei Wash Stock.

#### Dextran Buffer (22.5% w/v)

Dissolve 2.25 g dextran in 7 mL DPBS (no calcium, no magnesium). Use an end-to-top shaker for dissolving. Add 100 µL PIC (100X) and adjust volume to 10 mL with DPBS (no calcium, no magnesium). Sterile filter and keep at 4°C for no longer than 1 week.

#### FANS Buffer

Add 750 µL 20% BSA, 150 µL PIC (100X) and 30 µL 0.5 M EDTA to 14.07 mL DPBS (no calcium, no magnesium).

#### 2x Tn5 Loading Buffer

Add 1 mL 1 M HEPES pH 7.2, 400 µL 5 M NaCl, 4 µL 0.5 M EDTA, 200 µL 10% (vol/vol) Triton X-100 stock solution, 200 µL 100% glycerol to 6.4 mL nuclease-free water. Store at 4°C for up to 6 months.

On the day of use, add 2 µL 0.1 M DTT to 98 µL 2x Tn5 Loading Buffer.

#### 2x nanoCUT&Tag Wash Buffer

Add 2 mL 1 M HEPES pH 7.5 and 3 mL 5 M NaCl to 45 mL nuclease-free water. Store at 4°C for up to 6 months.

#### 5% DIG

Dissolve 100 mg Digitonin in 2 mL DMSO. Store at −20°C for up to 6 months.

! CAUTION Digitonin is toxic if swallowed, toxic in contact with skin, and fatal if inhaled. Causes skin and serious eye irritation; may cause respiratory tract irritation. Handle only with gloves, eye protection, face protection, and when dust/mist exposure is possible, use suitable respiratory protection. Ensure good ventilation.

#### 10% (vol/vol) NP-40 (IGEPAL CA-630)

Add 1 mL NP-40 (IGEPAL CA-630) to 9 mL nuclease-free water. Store protected from light at 20-22°C for up to 6 months.

#### 1 M MnCl_2_

Dissolve 0.1979 mg in 1 mL nuclease-free water. Store at 20-22°C for up to 6 months.

! CAUTION Manganese(II) chloride is harmful if swallowed or inhaled; causes skin and serious eye irritation; may cause respiratory irritation. Chronic inhalation or repeated exposure can cause neurological effects (manganism). Handle with gloves, eye/face protection, and avoid creating dust or aerosols.

#### 2 M Spermidine

Add 62.5 µL spermidine to 137.5 µL nuclease-free water. Store at −20°C for up to 6 months.

! CAUTION Spermidine solutions can cause skin and eye irritation and may cause mild respiratory irritation if aerosolised. Handle with gloves and eye protection. Avoid splashing or generating aerosols. Work in a well-ventilated area or fume hood and wash hands after handling.

#### Sort Buffer (3 rxns)

Add 1.125 µL 2 M Spermidine, 45 µL PIC (100X) and 2204 µL nuclease-free water to 2250 µL 2x nanoCUT&Tag Wash Buffer. Keep at 4°C and use within a week.

#### DIG-300 Buffer (3 rxns)

Add 300 µL 5 M NaCl, 2.5 µL 2 M spermidine, 20 µL 5% DIG, 100 µL PIC (100X), 10 µL 10% (vol/vol) NP-40 and 4568 µL nuclease-free water to 5000 µL 2x nanoCUT&Tag Wash Buffer. Use within a week

#### Antibody Wash Buffer (3 rxns)

Add 2.5 µL 2 M spermidine, 100 µL 5% DIG, 100 µL PIC (100X), 20 µL 10% (vol/vol) NP-40 and 4788 µL nuclease-free water to 5000 µL 2x nanoCUT&Tag Wash Buffer. Keep at 4°C and use within a week.

#### Complete Antibody Buffer (3 rxns)

Prepare on the day of the experiment and keep at 4°C. Take 400 µL of Antibody Wash Buffer and add 2 µL 20% BSA and 1.6 µL 0.5 M EDTA.

#### Binding Buffer

Add 1 mL 1 M HEPES pH 7.9, 500 µL 1 M KCl, 50 µL 1 M CaCl_2_, 50 µL 1 M MnCl_2_ to 49.4 mL nuclease-free water. Store at 4°C for up to 6 months.

#### Tagmentation Buffer (3 rxns)

Add 5 µL 1 M MgCl_2_ to 495 µL DIG-300 Buffer. Keep at 4°C and use within a week.

#### 2x TD Buffer

Add 40 µL 1 M HEPES pH 7.5, 20 µL 1M MgCl_2_ to 940 µL nuclease-free water. Store at 4°C for up to 6 months.

#### Oligonucleotides

Reconstitute the MeA, MeB, MeRev and indexing PCR primer oligonucleotides in water to a final concentration of 100 µM each. Keep these as a stock at −20 °C. Dilute an aliquot of the indexing PRC primers to a working concentration of 10 µM.

### Procedure

#### Preparation of reagents and equipment (Timing ∼15 min)

1. Per sample, prepare 20 mL Nuclei Wash Buffer in a 50 mL tube, 5 mL Lysis Buffer in a 15 mL tube, 10 mL Dextran Buffer in a 15 mL tube, 1.5 mL Sort Buffer in a 2 mL tube and 15 mL FANS buffer in a 15 mL tube fresh on the day of use as described in the “Reagents setup” section. Pre-cool all reagents on ice.
2. Activate biosafety cabinet airflow for homogenization of human tissue.
3. Set centrifuges to 4°C.

#### Tissue preparation (Timing ∼15 min)

! CRITICAL Do not thaw the brain tissue until step 8.

4. Pre-cool the forceps, razor blade, and petri dish on dry ice.
5. Transfer the frozen brain tissue from its storage container onto the petri dish on dry ice using the pre-cooled forceps. Avoid direct contact of the tissue with your hands or warm surfaces.

! CAUTION Human tissue must be treated as potentially pathogenic. Always handle human tissue using appropriate personal protective equipment (PPE) inside a Class II A2 biosafety cabinet, following institutional safety guidelines.

6. Cut ∼250 mg of brain tissue using the pre-cooled razor blade.
7. Prepare a petri dish on ice and add 5 mL ice-cold Lysis Buffer.
8. Transfer the tissue onto the petri dish with Lysis Buffer on ice using the pre-cooled forceps.
9. Thaw the brain tissue on ice for ∼5 min.

! CRITICAL: For tissue dissociation (steps 10-33), do not exceed ∼500 mg of tissue to avoid material loss during homogenization.

? Troubleshooting

#### Tissue dissociation (Timing ∼30 min)

10. Carefully mince the thawed brain tissue into ∼1 mm chunks using the precooled razor blade.
11. Transfer the minced brain tissue into a 7 mL Dounce using a 1 mL wide-bore pipette tip.
12. Let brain tissue pieces sink and use supernatant to flush the petri dish and transfer remaining tissue pieces to the Dounce.
13. Insert the loose pestle and slowly move it up and down 15 times without creating bubbles.
14. Insert the tight pestle and slowly move it up and down 10 times without creating bubbles.

! CRITICAL: Brain tissue homogenization should be gentle, avoiding excessive force and speed. Avoid creating bubbles. When the pestle is removed, ∼500 µL Nuclei Wash Buffer can be used to rinse the pestle.

15. Place a 70 µm strainer on top of a 50 mL tube and place on ice.
16. Pass the homogenate through the 70 µm strainer into the 50 mL tube.
17. Gently mash tissue pieces remaining on the 70 µm strainer using the rubber head of a 5 mL syringe.
18. Flush the strainer with 2 mL Nuclei Wash Buffer.
19. Repeat Steps 17-18.
20. Place a 40 µm strainer on top of a new 50 mL tube and place on ice.
21. Pass the homogenate through the 40 µm strainer into the 50 mL tube.
22. Gently mash the remaining tissue pieces on the 40 µm strainer using the rubber head of a 5 mL syringe.
23. Flush the strainer with 2 mL Nuclei Wash Buffer.
24. Repeat Steps 22-23.
25. Transfer the homogenate to a new 15 mL tube.
26. Rinse the 50 mL tube with 2 mL Nuclei Wash Buffer and transfer this to the 15 mL tube (total volume ∼15 mL).

#### Isolation of nuclei (Timing ∼45 min)

27. Spin the homogenate at 600xg for 10 min at 4°C.

! CRITICAL: Dextran Buffer is recommended for myelin removal. For samples that do not contain high amounts of myelin, proceed directly to step 31 and repeat the wash steps 31-32 one more time.

28. Remove the supernatant and resuspend the pellet in 2 mL Sort Buffer.
29. Add 8 mL Dextran Buffer to the homogenate and mix well by gently inverting 10 times.
30. Set the centrifuge deceleration to 6 and spin the homogenate at 4300xg for 15 min at 4°C.

? Troubleshooting

! CRITICAL: If no pellet is visible after step 30, repeat the centrifugation with the same settings for another 5-10 min.

31. Remove the supernatant and resuspend the pellet in 5 mL Nuclei Wash Buffer.
32. Spin the homogenate at 600xg for 10 min at 4°C.
33. Remove the supernatant and resuspend the pellet in 600 µL FANS Buffer.

#### Primary antibody staining for cell type enrichment of nuclei by FANS (Timing ∼16 hr)

34. For single stain controls of each antibody and for the DAPI unstained control, prepare FACS tubes with 0.3 mL FANS Buffer and add 3 µL of nuclei (from Step 33).
35. For single-stain controls, add the concentration of antibody indicated in Table 1 and protect from light using aluminium foil.
36. The remaining nuclei from step 35 are used for total stain (∼0.6 mL). Add the concentration of antibody indicated in Table 1 and protect from light using aluminium foil.
37. Mix by gently shaking the tubes for 10 seconds.
38. Incubate the nuclei suspension for 16 hours at 4°C.
39. The next day, add 3.5 mL FANS Buffer to each tube.
40. Spin the total stain and single stain tubes at 600xg for 10 min at 4°C.
41. Carefully discard the supernatant without disturbing the nuclei pellet.
42. Resuspend the unstained and single stain controls in 0.3 mL FANS Buffer. Resuspend the total stain sample in 1 mL FANS Buffer. Resuspend nuclei by gently pipetting up and down ten times.
43. Prepare 35 µm cell strainer snap caps by wetting the membrane with DPBS.
44. Transfer total stain and single stain samples through the 35 µm cell strainers.
45. Wash the total stain tube with 0.5 mL FANS Buffer and pass this through the 35 µm cell strainer (total volume 1.5 mL).
46. Add DAPI and incubate for 30 min (2 µL 1 mg mL^-1^ DAPI for total stain, and 1 µL 0.1 mg mL^-1^ DAPI for single stain and unstained controls).

#### Secondary antibody staining for unconjugated primary antibodies (Timing ∼45 min)

47. After step 42, add secondary antibodies according to Table 1 to the total stains, as well as the single stains for LHX2, RFX4 and SUR2B.
48. Incubate at 4°C for 1 hour.
49. Add 3.5 mL FANS Buffer to each tube.
50. Spin the total stain and single stain tubes at 600xg for 10 min at 4°C.
51. Carefully discard the supernatant without disturbing the nuclei pellet.
52. Resuspend unstained and single stain controls in 0.3 mL FANS Buffer and resuspend the total stain sample in 1 mL FANS Buffer by gently pipetting up and down ten times.
53. Prepare 35 µm cell strainer snap caps by wetting the membranes with DPBS.
54. Transfer total stain and single stain samples through the 35 µm strainer.
55. Wash the total stain tube with 0.5 mL FANS Buffer and pass this through the 35 µm cell strainer (total volume 1.5 mL).
56. Add DAPI and incubate for 30 min (2 µL 1 mg mL^-1^ DAPI for total stain, and 1 µL 0.1 mg mL^-1^ DAPI for single stain and unstained controls).

! Pause point Recommended cell type combinations for neurons (NeuN-488), microglia/myeloid cells (PU1-PE), and brain endothelial cells (ERG/FLI1-647). A subset of myeloid cells will be positive for ERG/FLI1; to enrich for brain endothelial cells, gate for the PU1^-ve^ and ERG/FLI1^high^ population. Recommended cell type combinations for human astrocytes (RFX4) or mouse astrocytes (LHX2) and neurons (NeuN-488); to remove contaminating glia, include a dump channel for oligodendrocytes (OLIG2-647) and microglia (PU1-647). Neurons express LHX2 and RFX4; to enrich for astrocytes, gate for the NeuN^-ve^ and LHX2^+ve^ (mouse) or RFX4^+ve^ (human) nuclei.

**Table 1.**

| Target | Clone | Conjugation | Host | Product conc. | Specificity | Company | Cat# | RRID | FANS conc. |
| --- | --- | --- | --- | --- | --- | --- | --- | --- | --- |
| anti-NeuN | EPR12763 | Alexa Fluor 488 | Rabbit | 0.5 mg mL<sup>-1</sup> | Human | Abcam | ab190195 | AB_2716282 | 1:2,500 |
| anti-NeuN | A60 | Alexa Fluor 488 | Mouse | 1.0 mg mL<sup>-1</sup> | Human, Mouse | SigmA-Aldrich | MAB377X | AB_2149209 | 1:2,500 |
| anti-PU1 | E8I8L | PE | Mouse | 0.1 mg mL<sup>-1</sup> | Human, Mouse | Cell Signaling | 20251 | AB_3716421 | 1:100 |
| anti-PU1 | E8I8L | Alexa Fluor 647 | Mouse | 0.1 mg mL<sup>-1</sup> | Human, Mouse | Cell Signaling | 20698 | AB_3716422 | 1:100 |
| anti-OLIG2 | EPR2673 | Alexa Fluor 488 | Rabbit | 0.5 mg mL<sup>-1</sup> | Human, Mouse | Abcam | ab225099 | AB_3714884 | 1:2,500 |
| anti-OLIG2 | EPR2673 | Alexa Fluor 647 | Rabbit | 0.5 mg mL<sup>-1</sup> | Human, Mouse | Abcam | ab225100 | AB_3675495 | 1:2,500 |
| anti-OLIG2 | OLIG2/2400 | Cyanine Fluor 647 | Mouse | 0.1 mg mL<sup>-1</sup> | Human, Mouse | Biotium | BNC472400 | AB_3716423 | 1:100 |
| anti-LHX2 | polyclonal | Unconjugated | Rabbit | 0.2 mg mL<sup>-1</sup> | Human | Abcam | ab219983 | AB_2868535 | 1:200 |
| anti-LHX2 | EPR20449 | Unconjugated | Rabbit | 0.5 mg mL<sup>-1</sup> | Mouse | Abcam | ab184337 | AB_2916270 | 1:200 |
| anti-RFX4 | polyclonal | Unconjugated | Rabbit | 0.2 mg mL<sup>-1</sup> | Human | Abcam | ab188020 | AB_3716420 | 1:200 |
| anti-ERG | EPR3864 | Alexa Fluor 488 | Rabbit | 0.5 mg mL<sup>-1</sup> | Human, Mouse | Abcam | ab196374 | AB_2889273 | 1:100 |
| anti-ERG | EPR3864 | Alexa Fluor 647 | Rabbit | 0.5 mg mL<sup>-1</sup> | Human, Mouse | Abcam | ab196149 | AB_3697463 | 1:100 |
| anti-NOTCH3 | MHN3-21 | PE | Mouse | 0.1 mg mL<sup>-1</sup> | Human | BioLegend | 345405 | AB_2151232 | 1:100 |
| anti-SUR2B | N323A/31 | Unconjugated | Mouse | 1.0 mg mL<sup>-1</sup> | Mouse | Thermo Fisher Scientific | 75-298 | AB_2315926 | 1:2500 |
| anti-Rabbit IgG | polyclonal | PE | Goat | 2.0 mg mL<sup>-1</sup> | Rabbit IgG | Cell Signaling | 79408 | AB_2799931 | 1:1,000 |
| anti-Mouse IgG | polyclonal | Alexa Fluor 647 | Donkey | 2.0 mg mL<sup>-1</sup> | Mouse IgG | Thermo Fisher Scientific | A32787 | AB_2762830 | 1:1,000 |

#### FANS of cell type-enriched nuclei (Timing ∼4 hr)

57. Count nuclei using a hemacytometer and dilute to approximately 1×10⁶ nuclei mL^-1^.

! CRITICAL Suspensions with a high nuclei concentration can clog the sorter. Debris from the isolation procedure can interfere with accurate nuclei counting. As a guide, nuclei isolated from ∼250 mg of brain tissue are typically diluted in ∼1.5 mL FANS Buffer for sorting. Take extra FANS buffer, collection tubes and DAPI to the sorter for further dilution, if needed.

58. Pre-fill 1.5 or 2 mL DNA low-Bind collection tubes with 500 µL of Sort Buffer.
59. Sort unfixed nuclei using either an 85 µm or 100 µm nozzle.

! Pause point An 85 µm nozzle provides higher sorting resolution and faster droplet generation for clean samples, but is more prone to clogging, particularly with debris-rich preparations. A 100 µm nozzle provides more stable sorting at higher flow rates. If clogging occurs, dilute the nuclei suspension with additional FANS Buffer.

60. Use unstained and single-stained controls to establish fluorescence compensation and assess spectral overlap before initiating the sort.
61. Using the total stained sample, generate a dot plot of forward scatter area (FSC-A; x-axis) versus DAPI (y-axis) to gate for DAPI^+ve^ nuclei (**Fig. 2A**).
62. From the DAPI^+ve^ population, generate a dot plot of FSC-A (x-axis) versus FSC-H (y-axis) to gate for singlet DAPI^+ve^ nuclei and to deplete aggregates (**Fig. 2A**).
63. To enrich for neuronal and myeloid/microglia nuclei, plot NeuN-488 versus PU1-PE from the singlet DAPI^+ve^ population and gate NeuN^+ve^ and PU1^+ve^ nuclei, respectively (Fig. 2B).
64. To enrich for brain endothelial nuclei, plot NeuN-488 versus ERG-647 from the NeuN^-ve^/PU1^-ve^ population and gate for ERG^high^ nuclei (**Fig. 2B**).
65. To enrich for astrocytes, plot NeuN-488 versus RFX4-PE (human) or LHX2-PE (mouse) from the NeuN^-ve^, OLIG2^-ve^/PU1^-ve^ population and gate for RFX4^+ve^ (human) nuclei^13^ or LHX2^+ve^ (mouse) nuclei (**Fig. 2C**).
66. To enrich for human mural cells, plot OLIG2-488 versus NOTCH3-PE from the OLIG2^-ve^ and PU1^-ve^ population and gate for NOTCH3^+ve^ nuclei^13^.
67. To enrich for mouse mural cells, plot ERG-488 versus SUR2B-PE from the ERG^-ve^ and PU1^-ve^ population and gate for SUR2B^+ve^ nuclei (**Fig. 2D**).
68. Cell-type-enriched nuclei are collected using high-purity sort mode into 20% BSA coated 1.5 mL tubes pre-filled with 500 µL Sort Buffer at 4 °C. Collect ∼10,000-50,000 nuclei for nanoCUT&Tag. Additional nuclei can be processed and stored (step 71).

! Pause point Assess nuclei purity by reanalyzing 5% of the sorted nuclei resuspended in 300 µL of FANS buffer in a new FACS tube and analyzed using the same sort parameters. For high-purity populations, >95% of nuclei should fall within the original sorting gates.

? Troubleshooting

#### Nuclei storage for downstream applications (Timing ∼20 min)

69. For nuclei storage, spin at 600xg for 10 min at 4 °C using a swinging-bucket rotor.
70. Carefully remove the supernatant and resuspend in 50 µL Bambanker per 50,000 nuclei.
71. Store the nuclei at −80 °C. Unfixed nuclei can be stored for several months in Bambanker at −80 °C without substantial loss of quality.

#### Loading Tn5 (Timing 1.5 hr)

72. Add 10 µL 100 µM MeA to 10 µL 100 µM MeRev (tube 1). And add 10 µL 100 µM MeB to 10 µL 100 µM MeRev (tube 2).
73. Denature the 50 µM MeA/MeRev and 50 µM MeB/MeRev oligonucleotides in a thermocycler at 95°C for 5 min and ramp down to 20°C by 0.1 °C per second. Annealed oligonucleotides can be stored at −20°C.
74. Mix 4 µL annealed 50 µM MeA/Me-Rev oligos with 21 µL glycerol, 1.5 µL (15 µg) mouse nanoTn5 or rabbit nanoTn5 and 23.5 µL of 2x Tn5 loading buffer (with DTT) (50 µL final volume).

<u>! CRITICAL</u> The volume of nanoTn5 needs to be adjusted according to the concentration of each Tn5 batch

75. Mix 4 µL annealed 50 µM MeB/Me-Rev oligos with 21 µL glycerol, 5 µL Tn5 Transposase - unloaded (Cat# C01070010) and 20 µL of 2x Tn5 loading buffer (with DTT).
76. Mix well by gently pipetting up and down ten times.
77. Incubate for 1 hour at 20-22°C (room temperature).
78. Store the loaded Tn5 at −20°C.

#### Concanavalin bead preparation (Timing ∼10 min)

79. Resuspend the Concanavalin A beads by pipette mixing or gentle inversion.
80. For each sample, add 20 µL of Concanavalin A beads to a 2 mL protein low-bind tube.
81. Add 1.6 mL of Binding Buffer to each 2 mL tube containing the Concanavalin A beads. Mix thoroughly by inversion or pipetting.
82. Place the 2 mL tube containing the bead slurry on a magnetic stand. Allow the solution to clear (∼2 min) and then remove the supernatant.
83. Resuspend beads in 1.5 mL Binding Buffer. Mix well by inversion or pipetting.
84. Return the tube to the magnetic stand. Allow the solution to clear (∼2 min) and then remove the supernatant.
85. Resuspend the beads in 20 µL Binding Buffer. Mix well by pipetting and keep on ice.

? Troubleshooting

#### nanoCUT&Tag (Day 1) (Timing ∼16 hr)

86. Briefly spin the 10,000-50,000 FANS-sorted nuclei from step 68.
87. Fill the tube with Sort Buffer to a final volume of 1.5 mL.
88. Add 1.5 mL nuclei (step 87) to 20 µL Concanavalin A beads (step 85).
89. Mix by inversion followed by 10 min rocking on an end-to-top rotator at 4^°^C for 10 min.
90. Spin briefly and place on a magnetic stand. Allow the solution to clear (∼2 min) and then remove the supernatant.
91. Resuspend the nuclei/beads in 50 µL Complete Antibody Buffer.
92. Pipette mix and keep on ice.
93. Add 1 µL histone antibody (**Table 2**) to each sample and mix by gently flicking the tube

**Table 2.**

| Target | Clone | Host | Product conc. | Specificity | Company | Cat# | RRID | nanoCUT&Tag conc. |
| --- | --- | --- | --- | --- | --- | --- | --- | --- |
| anti-H3K27ac | polyclonal | Rabbit | 1.0 mg mL<sup>-1</sup> | Human, Mouse, Rat, Cow | Abcam | ab4729 | AB_2118291 | 1:50 (1µg) |
| anti-H3K27ac | monoclonal | Mouse | 1.0 mg mL<sup>-1</sup> | Human, Mouse, Rat | Thermo Fisher Scientific | MA523516 | AB_2608307 | 1:50 (1µg) |
| anti-H3K4me3 | monoclonal (MC315) | Rabbit | NA | Human | Merck Millipore | 04-745 | AB_1163444 | 1:50 |
| anti-H3K4me3 | monoclonal | Mouse | 1.0 mg mL<sup>-1</sup> | Human, Mouse, Nematodes, Arabidopsis | Diagenode | C15200152 | AB_3716424 | 1:50 (1µg) |
| anti-H3K27me3 | monoclonal (C36B11) | Rabbit | NA | Human, Mouse, Rat, Monkey | Cell Signaling | 9733 | AB_2616029 | 1:50 |
| anti-H3K27me3 | monoclonal | Mouse | 1.0 mg mL<sup>-1</sup> | Human, Nematodes, Magnaporthe oryzae | Diagenode | C15200181 | AB_2819192 | 1:50 (1µg) |

! CRITICAL Host species of the primary histone antibody should differ from the antibody used for FANS.

94. Incubate for 16 hours at 4°C on an orbital shaker.

? Troubleshooting

#### First tagmentation and clean up (Timing ∼ 4.5 hr)

95. The next day, spin briefly and place on a magnetic stand. Allow the solution to clear (∼2 min) and then remove the supernatant.
96. Add 1 mL Antibody Wash Buffer and gently shake or slowly pipette mix up and down.
97. Spin briefly and place on a magnetic stand. Allow the solution to clear (∼2 min) and then remove the supernatant.
98. Repeat wash steps 96-97 two times with Antibody Wash Buffer for a total of three washes.
99. Resuspend the nuclei/beads mix in 50 µL Complete Antibody Buffer.
100. Add 0.5 µL nanoTn5 loaded with Me-A/Me-Rev (step 74) that matches the antibody species for the targeted histone (mouse or rabbit).
101. Mix by gently shaking the tube and place on an orbital shaker or thermomixer at 350 rotations per minute (rpm) and incubate for 1 hour at 20-22°C.
102. Spin briefly and place on a magnetic stand. Allow the solution to clear (∼2 min) and then remove the supernatant.
103. Add 1 mL Dig-300 Buffer and gently shake or slowly pipette mix.
104. Spin briefly and place on a magnetic stand. Allow the solution to clear (∼2 min) and then remove the supernatant.
105. Repeat steps 103-104 two more times with Dig-300 Buffer for a total of three washes.
106. Add 125 µL Tagmentation Buffer and gently shake or slowly pipette mix.
107. Incubate in a thermomixer for 60 min at 37°C at 350 rpm.
108. Remove samples and spin briefly. Set the temperature of the thermomixer to 55°C.
109. To stop tagmentation and solubilise DNA, add 4.2 µL 0.5 M EDTA, 1.25 µL 10% SDS and 1.1 µL 10 mg mL^-1^ Proteinase K.
110. Mix by vortexing at full speed for ∼2 seconds and incubate in a thermomixer for 60 min at 55°C at 350 rpm.
111. Spin briefly and place on a magnetic stand. Allow the solution to clear (∼2 min) and transfer ∼135 µL of the sample to a new 1.5 mL DNA low-bind tube.
112. Perform the following steps at 20-22°C. Add 675 µL Zymo DNA Binding Buffer to 135 µL sample (step 111) and invert mix.
113. Transfer the sample to a Zymo Spin Column and place into a Collection Tube.
114. Spin at 10,000xg for 30 seconds and discard the flow-through.
115. Add 200 µL Zymo Wash Buffer (with ethanol) to the Zymo Spin Column.
116. Spin at 10,000xg for 30 seconds and discard the flow-through.
117. Repeat steps 115-116.
118. Dry the Zymo Spin Column by spinning at 10,000xg for 1 min.
119. Place the Zymo Spin Column in a 1.5 mL DNA low-bind tube and add 24 µL of nuclease-free water. Incubate for 1 minute at 20-22°C.
120. Spin at 10,000xg for 1 minute. Discard the Zymo Spin Column and keep the eluate on ice.

! Pause point The sample can be stored at −20°C for 2 weeks.

#### Linear PCR and clean up (Day 3) (Timing: ∼45 min)

121. Mix 23 µL tagmented DNA (from step 120) with 2 µL 10 µM i5 primer and 25 µL NEBNext Ultra II Q5 MM.
122. Run samples on the following PCR steps. Step 1: 72°C for 5 min; Step 2: 98°C for 30 sec; Step 3: 98°C for 10 sec; Step 4: 63°C for 10 sec; Step 5: 72°C for 1 min; Step 6: 10°C hold. Repeat PCR steps 3-4 for 7 to 10 cycles. Cycle number should be optimised for the number of input nuclei and the nanoCUT&Tag target antibody.
123. Prepare 500 µL 80% ethanol per sample. Keep at 20-22°C.
124. Mix SPRI beads thoroughly by pipetting up and down.
125. Add 55 µL SPRI beads to the sample (1.1x) and mix well by pipetting at 20-22°C.
126. Incubate for 5 min at 20-22°C.
127. Spin briefly and place on a magnetic stand. Allow the solution to clear (∼2 min) and remove the supernatant.
128. Keep on the magnetic stand, add 200 µL 80 % ethanol and incubate for 30 sec at 20-22°C.
129. Remove the ethanol wash and repeat step 128.
130. Spin briefly and place on a magnetic stand. Wait for 30 seconds and discard residual ethanol.
131. Keep the tube open at 20-22°C and wait 2-5 minutes until the beads become matte.
132. Remove from the magnetic stand and resuspend beads with 50 µL nuclease-free water.
133. Incubate for 5 minutes at 20-22°C.
134. Spin briefly and place on a magnetic stand. Allow the solution to clear (∼2 min).
135. Transfer 50 µL DNA sample to a new 2 mL DNA low-bind tube.

! Pause point Sample can be stored at −20°C for 2 weeks.

#### Second tagmentation and clean up (Timing ∼45 min)

136. Quantify DNA using Qubit™ 1X dsDNA HS Assay or equivalent using manufacturer’s instructions.
137. Transfer 5 ng DNA to a new 2 mL DNA low-bind tube and adjust volume to 50 µL with nuclease-free water.
138. Add 50 µL 2x TD buffer to 50 µL 5 ng DNA sample.
139. Add 0.2 µL MeB/MeRev loaded Tagmentase.
140. Incubate in a thermomixer set at 37°C for 30 min at 350 rpm.
141. Remove samples from the thermomixer and directly proceed to DNA clean up.
142. Perform the following steps at 20-22°C. Add 500 µL Zymo DNA Binding Buffer to 100 µL sample (step 140) and invert mix.
143. Transfer the sample to a Zymo Spin Column and place into a Collection Tube.
144. Spin at 10,000xg for 30 seconds and discard the flow-through.
145. Add 200 µL Zymo Wash Buffer (with ethanol) to the Zymo Spin Column.
146. Spin at 10,000xg for 30 seconds and discard the flow-through.
147. Repeat steps 145-146.
148. Dry the Zymo Spin Column by spinning at 10,000xg for 1 min.
149. Place the Zymo Spin Column in a 1.5 mL tube and add 22 µL of nuclease-free water. Incubate for 1 minute at 20-22°C.
150. Spin at 10,000xg for 1 minute. Discard the Zymo Spin Column and keep the elute in the 1.5 mL tube on ice.

Safe point: The sample can be stored at −20°C for 2 weeks.

#### Final exponential PCR and clean up (Timing ∼45 min)

151. Add 21 µL DNA (step 150) to 2 µL 10 µM i5 primer, 2 µL 10 µM i7 primer and 25 µL NEBNext Ultra II Q5 MM and pipette mix.
152. Run samples on the following PCR steps. Step 1: 72°C for 5 min; Step 2: 98°C for 30 sec; Step 3: 98°C for 10 sec; Step 4: 63°C for 10 sec; Step 5: 72°C for 1 min; Step 6: 10°C hold. Repeat PCR steps 3-4 for 7 to 10 total cycles. Cycle number should be optimised for the number of input nuclei and the nanoCUT&Tag target antibody.
153. Prepare 500 µL 80% ethanol per sample. Keep at 20-22°C.
154. Mix SPRI beads thoroughly by pipetting up and down.
155. Add 55 µL SPRI beads to the sample (1.1x) and mix well by pipetting at 20-22°C.
156. Incubate for 5 min at 20-22°C.
157. Spin briefly and place on a magnetic stand. Allow the solution to clear (∼2 min) and then remove the supernatant.
158. Keep on the magnetic stand, add 200 µL 80 % ethanol and incubate for 30 sec at 20-22°C.
159. Remove the ethanol wash and repeat step 158.
160. Spin briefly and place on a magnetic stand. Wait for 30 seconds and discard residual ethanol.
161. Keep the tube open at 20-22°C and wait 2-5 minutes until the beads become matte.
162. Remove from the magnetic stand and resuspend beads with 20 µL nuclease-free water.
163. Incubate for 5 minutes at 20-22°C.
164. Spin briefly and place on a magnetic stand. Allow the solution to clear (∼2 min).
165. Transfer 20 µL DNA sample to a new 1.5 mL DNA low-bind tube.

Safe point: The sample can be stored at −20°C or −80°C for several years.

#### Library submission (Timing: ∼30 min)

166. Quantify DNA using Qubit™ 1X dsDNA HS Assay or equivalent using manufacturer’s instructions.
167. Assess DNA size distribution using an Agilent 2100 Bioanalyzer and Agilent High Sensitivity DNA Kit, or Agilent TapeStation and D5000 High Sensitivity ScreenTape Assay, or equivalent. Optimal average fragment size is 400-600 bp.
168. Adjust the DNA library volume and concentration for next-generation sequencing according to the platform or facility requirements.

? Troubleshooting

### Troubleshooting

**Step:** 9

**Problem:** Deterioration of tissue quality due to repeated removal from the −80°C freezer.

**Possible reason:** The block of tissue is larger than needed per experiment. Removing the tissue from the freezer multiple times to cut an aliquot causes repeated partial thawing, causing deterioration in tissue quality.

**Possible solution:** For larger blocks of tissue, aliquot these into separate containers containing the amount needed for one experiment to avoid repeated removal of the sample from the freezer.

**Step:** 30

**Problem:** No clear nuclei pellet is visible after centrifugation.

**Possible reason:** The nuclei homogenate was not properly mixed with the dextran.

**Possible solution:** Homogenise the sample and dextran again by inverting at least 10 times. The sample should be completely homogenous, and then the centrifugation step can be repeated.

**Step:** 61 - 68

**Problem:** High background fluorescence makes it difficult to identify false-positive nuclei.

**Possible reason:** Autofluorescence is a common issue in human brain samples, due to the handling of the tissue and longer post-mortem intervals.

**Possible solution:** High-quality tissue, with a lower post-mortem interval, should be selected. During FANS, nuclei of interest should be gated using two different fluorescent channels. When plotting two channels on one plot, auto-fluorescent nuclei will be double-positive and can be avoided when gating the nuclei of interest.

**Step:** 61 - 68

**Problem:** A lower-than-expected number of nuclei per cell-type during FANS.

**Possible reason:** Antibody staining is suboptimal. Myelin present in the sample will bind and deplete antibodies.

**Possible solution:** Resuspend the nuclei pellet in a larger volume of FANS buffer or increase the amount of antibody used.

**Step:** 61 - 68

**Problem:** The cell type of interest is isolated by staining for two or more nuclear markers using antibodies raised in both mouse and rabbit, which prevents species-specific CUT&Tag analysis using nanoCUT&Tag.

**Possible reason:** If nuclei are stained for two or more transcription factors that are raised in both mouse and rabbit, the mouse and rabbit nanobody-Tn5 will bind unspecifically to these transcription factors.

**Possible solution:** When using two or more nuclear markers that stain the same cell type, these should be raised in the same host or not raised in rabbit and mouse to be compatible with the mouse and rabbit nanobody-Tn5. Alternatively, it has been reported that antibodies used in FANS can be blocked using fragment-antigen binding (Fab) fragments^25^, avoiding the need for species-specific nano-Tn5 for the CUT&Tag protocol.

**Step:** 68

**Problem:** FANS-enrichment of cell type nuclei does not reach the minimum recommended number of 10,000-50,000 nuclei needed for nanoCUT&Tag.

**Possible reason**: The cell type of interest is rare, making it difficult to reach the minimum number of nuclei needed for nanoCUT&Tag

**Possible solution:** nanoCUT&Tag is optimised for 10,000 – 50,000 nuclei. Lower numbers will impact the data quality. For samples with <10,000 nuclei, it is still possible to proceed with the nanoCUT&Tag, however, this may impact data quality.

**Step:** 79 - 85

**Problem:** The concanavalin A bead slurry is sticking to the tube walls, causing them to dry out.

**Possible reason:** The bead slurry is more prone to stick to certain types of plastic.

**Possible solution:** It is important to use protein low-bind tubes for steps of the protocol where the nuclei are intact. Once the DNA has been extracted from the nuclei, it is important to use DNA low-bind tubes to prevent DNA loss.

**Step:** 86 - 94

**Problem:** Concanavalin A beads are becoming sticky, making it difficult to remove the supernatant

**Possible reason:** Nuclei are lysing too early, and DNA leaks out, causing the nuclei and beads to become sticky.

**Possible solution:** The amount of digitonin and NP-40 can be reduced, if necessary. Note that the purity of the digitonin used in this protocol is 50% (TLC).

**Step:** 166

**Problem:** Sample concentration is lower than the sequencing facility’s requirements.

**Possible reason:** Some nanoCUT&Tag libraries have a lower library yield, such as nanoCUT&Tag for transcription factors and low-abundance histone modifications.

**Possible solution:** Cycle numbers per library type need to be optimised. Additional PCR cycles can be performed after quantifying the final library. If necessary, the final library can be eluted in a lower volume of nuclease-free water, or the sample volume can be reduced using a vacuum pump.

**Step:** 167

**Problem:** The sample is under- or over-tagmented, leading to sub-optimal sequencing conditions.

**Possible reason:** If samples are under- or over-tagmented, Tn5 should be titrated to the right concentration.

**Possible solution:** If less than 5 ng of DNA is carried over to tagmentation #2 (step 137), cycle numbers of the linear PCR can be increased. Tn5 tagmentase is extremely temperature sensitive. Minimise the time the Tn5 tagmentase is outside the −20°C freezer and keep in a benchtop mini cooler or on ice when in use.

### Anticipated results

The unfixed nuclei isolation followed by FANS-enrichment has routinely been used to isolate nuclei from defined cell populations from C57BL/6 mouse and human brain tissue. From ∼100 mg of mouse brain tissue, average nuclei yields were ∼108,000 microglia/myeloid cells, ∼18,300 brain endothelial cells, ∼12,000 mural cells, ∼1.65 million neurons, ∼495,000 oligodendrocytes, and ∼135,000 astrocytes. From human postmortem brains, 100 mg tissue yielded ∼60,000 microglia/myeloid cells, ∼8,900 BECs, ∼9,200 mural cells, ∼114,000 neurons, ∼165,000 oligodendrocytes, and ∼64,000 astrocytes. Slight differences were observed in human resected brain tissue, where 100 mg resulted in ∼168,000 microglia/myeloid cells, ∼4,300 brain endothelial cells, ∼3,600 mural cells, ∼269,000 neurons, ∼242,000 oligodendrocytes, and ∼25,000 astrocytes. It was previously observed that PU1 staining of microglia/myeloid cell nuclei in post-mortem tissue was often compromised, likely due to tissue quality^16^. This has been improved by a new PU1 antibody (Cell Signaling, Cat # 20251). Strong and consistent PU1 staining was detected using both resected and postmortem samples, with postmortem intervals tested up to ∼45 hours. Brain endothelial cells and mural cells were challenging to isolate, and optimal recovery required processing >250 mg human brain tissue to isolate ∼10,000 nuclei.

Profiling of histone modifications and transcription factors using nanoCUT&Tag was optimally performed using 10,000-50,000 nuclei, with good quality data still achievable with 5,000-10,000 nuclei. DNA concentrations after linear amplification ranged from 0.1 to 0.5 ng µL^-1^. The final library amplification resulted in DNA concentrations ranging from 1 to 10 ng µL^-1^. If higher DNA concentrations are consistently obtained after the linear amplification or final PCR, it is recommended to reduce the cycle number to minimise duplication rates. The fragment size distribution should display a unimodal size distribution centred between 400 and 600 bp (see troubleshooting for under- or over-tagmentation).

The protocol has been tested for multiple histone modifications, including H3K27ac (active gene regulatory elements), H3K4me3 (promoters), and H3K27me3 (repressed regions), which demonstrated specificity across cell types (**Fig. 3A**). While the protocol is compatible with some transcription factors, including ERG/FLI1, PU1, and OLIG2^13^, the high-salt concentrations used for nanoCUT&Tag may displace transcription factors with low DNA-binding affinities. The protocol is compatible with histone modifications across cell types using both mouse (**Fig. 3B**) and human post-mortem and resected brain tissue^13^. The nanoCUT&Tag samples for histone modifications were sequenced to ∼15 million read pairs with alignment rates >90%, fraction of reads in peaks (FRiP) >15%, duplication rates <45%, and linear duplication rates <55%^13^. nanoCUT&Tag samples for transcription factors had lower FRiP scores, which likely reflects an expected lower genome coverage compared to histone modifications. Overall, the protocol generated consistent high-quality nanoCUT&Tag datasets for histone modifications and some transcription factors.

The unfixed nuclei isolation is compatible with snRNA-seq, snNanoCUT&Tag and multiome snNanoCUT&Tag (H3K27ac+RNA). Approximately >30,000 unfixed FANS nuclei were required for snRNA-seq, and >200,000 FANS nuclei were required for snNanoCUT&Tag and multiome snNanoCUT&Tag experiments due to increased nuclei clumping. To enrich for vascular cell types, neurons, oligodendrocytes and microglia were depleted, which yielded ∼54,000 nuclei from ∼100 mg human brain tissue. Single-cell gene expression analysis indicated a ∼25% enrichment of brain endothelial cells and mural cells following depletion of neurons, oligodendrocytes and microglia (**Fig. 3C**).

For snNanoCUT&Tag, DNA concentrations for the final libraries were ∼1-10 ng µL^-1^, with expected fragment size distributions. H3K27ac snNanoCUT&Tag data is sparser compared to single-cell gene expression, as expected for single-cell epigenomic approaches. However, H3K27ac snNanoCUT&Tag analysis identified major cell type clusters, including vascular-associated brain endothelial cells, mural cells and astrocytes (**Fig. 3D**). To improve cell type clustering and annotation, multiome snNanoCUT&Tag (H3K27ac+RNA) was used to profile gene expression and H3K27ac in the same nucleus (**Fig. 3E, F**). The gene expression analysis for the multiome snNanoCUT&Tag dataset was used to annotate the H3K27ac clusters for microglia, neurons, oligodendrocytes and astrocytes (**Fig. 3E, F**). Overall, the multiome snNanoCUT&Tag approach produced complementary data that enhanced cell type resolution and annotation.

## Data availability

Human postmortem data was deposited at Gene Expression Omnibus under accession GSE299072^13^. Human resected data was deposited at EGA under accession EGAS50000001160^13^. Mouse data has been available at Gene Expression Omnibus under accession TBD.

## Acknowledgements

A.N. is supported by the UK Dementia Research Institute (UKDRI-5208) through UK DRI Ltd, principally funded by the Medical Research Council, The Dunhill Medical Trust (AISRPG2305\26), and the Alzheimer’s Association (ADSF-24-1345198-C). We thank Dr. Nyman, Dr. Strandback, Dr. Ampah-Korsah, and the Karolinska Institute Protein Science facility for nanobody-Tn5 production. The Imperial BRC Genomics Facility provided resources and support and is supported by NIHR funding to the Imperial Biomedical Research Centre. This work was supported by the Imperial College London Hammersmith flow cytometry facility, which is infrastructure funded by the NIHR Imperial Biomedical Research Centre (BRC); and the LMS/NIHR Flow Facility at Imperial.

## Contributions

K.C.Z., J.D.v.D., and A.N. conceptualized the study. K.C.Z., and J.D.v.D. optimized the FANS and nanoCUT&Tag methodology. L.B., and J.L.T. provided input into the FANS optimization. K.C.Z., and J.D.v.D. acquired the FANS and nanoCUT&Tag data. K.C.Z. analyzed the CUT&Tag datasets. K.C.Z., J.D.v.D., and A.N. wrote the manuscript.

## Key references using this protocol

Ziegler, K. C. *et al.* The brain neurovascular epigenome and its association with dementia. *Neuron* **114**, 268-286.e269, doi:10.1016/j.neuron.2025.10.001 (2026).

Nott, A., Schlachetzki, J. C. M., Fixsen, B. R. & Glass, C. K. Nuclei isolation of multiple brain cell types for omics interrogation. Nat Protoc 16, 1629-1646, doi:10.1038/s41596-020-00472-3 (2021).

Bárcenas-Walls, J. R. *et al.* Nano-CUT&Tag for multimodal chromatin profiling at single-cell resolution. *Nat Protoc* **19**, 791-830, doi:10.1038/s41596-023-00932-6 (2024).

## Notes

https://www.cell.com/neuron/fulltext/S0896-6273(25)00754-8

### Competing Interest Statement

The authors have declared no competing interest.

